# Isolation of cross-reactive monoclonal antibodies against divergent human coronaviruses that delineate a conserved and vulnerable site on the spike protein

**DOI:** 10.1101/2020.10.20.346916

**Authors:** Chunyan Wang, Rien van Haperen, Javier Gutiérrez-Álvarez, Wentao Li, Nisreen M.A. Okba, Irina Albulescu, Ivy Widjaja, Brenda van Dieren, Raul Fernandez-Delgado, Isabel Sola, Daniel L. Hurdiss, Olalekan Daramola, Frank Grosveld, Frank J.M. van Kuppeveld, Bart L. Haagmans, Luis Enjuanes, Dubravka Drabek, Berend-Jan Bosch

**Affiliations:** Division of Infectious Diseases and Immunology, Department of Biomolecular Health Sciences, Faculty of Veterinary Medicine, Utrecht University, Utrecht, the Netherlands; Department of Cell Biology, Erasmus Medical Center, Rotterdam, the Netherlands; Harbour BioMed, Rotterdam, the Netherlands; Department of Molecular and Cell Biology, National Center for Biotechnology-Spanish National Research Council (CNB-CSIC), Madrid, Spain; Department of Viroscience, Erasmus Medical Center, Rotterdam, the Netherlands; Cell Culture and Fermentation Sciences, Biopharmaceutical Development, BioPharmaceuticals R&D, AstraZeneca, Cambridge, United Kingdom

## Abstract

The coronavirus spike glycoprotein, located on the virion surface, is the key mediator of cell entry. As such, it is an attractive target for the development of protective antibodies and vaccines. Here we describe two human monoclonal antibodies, 1.6C7 and 28D9, that display a remarkable cross-reactivity against distinct species from three *Betacoronavirus* subgenera, capable of binding the spike proteins of SARS-CoV and SARS-CoV-2, MERS-CoV and the endemic human coronavirus HCoV-OC43. Both antibodies, derived from immunized transgenic mice carrying a human immunoglobulin repertoire, blocked MERS-CoV infection in cells, whereas 28D9 also showed weak cross-neutralizing potential against HCoV-OC43, SARS-CoV and SARS-CoV-2 in a neutralization-sensitive virus pseudotyping system, but not against authentic virus. Both cross-reactive monoclonal antibodies were found to target the stem helix in the spike protein S2 fusion subunit which, in the prefusion conformation of trimeric spike, forms a surface exposed membrane-proximal helical bundle, that is antibody-accessible. We demonstrate that administration of these antibodies in mice protects from a lethal MERS-CoV challenge in both prophylactic and/or therapeutic models. Collectively, these antibodies delineate a conserved, immunogenic and vulnerabe site on the spike protein which spurs the development of broad-range diagnostic, preventive and therapeutic measures against coronaviruses.

## Introduction

The Coronavrius Disease 2019 (COVID-19) pandemic is caused by the severe acute respiratory syndrome coronavirus 2 (SARS-CoV-2) which emerged in China, in late 2019 ^1^. The virus belongs to the subgenus *Sarbecovirus* of the genus *Betacoronavirus* within the subfamily *Orthocoronavirinae* ^2^. Two other zoonotic coronaviruses emerged in the last 20 years and cause severe acute respiratory disease in humans, similar to that seen with SARS-CoV-2. SARS-CoV, which also belongs to the subgenus *Sarbecovirus* and is closely related to SARS-CoV-2, crossed species barriers to humans in China in 2002, and caused worldwide ~8000 cases with a 10% mortality rate before it was contained in 2003 ^3^. Ten years later in 2012, the Middle East respiratory syndrome (MERS) coronavirus (subgenus *Merbecovirus*, genus *Betacoronavirus*) surfaced in Saudi Arabia. This virus is recurrently introduced in the human population from a dromedary camel reservoir with limited human-to-human spread and has led to ~2500 cases with approximately 35% of reported patients succumbing to the infection ^4^. In addition, four endemic human coronaviruses (HCoVs) circulate in humans, including HCoV-229E and HCoV-NL63 (genus *Alphacoronavirus*) and HCoV-OC43 and HCoV-HKU1 (subgenus *Embecovirus*, genus *Betacoronavirus*). While these viruses are typically associated with mild respiratory illnesses (common colds) ^5–7^, they can cause significant morbitity, and even mortality, in immunocompromised individuals ^8^. All these viruses made their way into humans from an animal reservoir, illustrating the zoonotic threat posed by coronaviruses and their pandemic potential ^1, 9–11^. The devastating socio-economic consequences of the COVID-19 pandemic urges the development of intervention strategies that can mitigate outbreak of future emerging coronaviruses.

Identification of antibodies that broadly react with existing (human) coronaviruses could be of relevance to increase our level of pandemic preparedness. Such antibodies might be useful for virus diagnostics in the early stages of a pandemic. Moreover, their epitopes may enable development of antibody-based therapeutics or vaccines that provide broad protection not only against contemporary pathogens but also against those that likely emerge in the future ^12^. Although being identified for other virus families ^13–18^, exploration of broadly reactive antibodies against coronaviruses is still in its infancy. Analysis of human polyclonal sera indicated that such cross-reactive antibodies might exist as one study reported that 25% of the convalescent SARS patients sera had low titers of antibodies that could neutralize MERS-CoV ^19^. In addition, Barnes *et al.* ^20^ reported reactivity of purified plasma IgG from ten convalescent COVID-19 patients with the MERS-CoV spike protein, and a study by Wec *et al.* ^21^ demonstrated that a subset of SARS-CoV-2 cross-reactive antibodies isolated from a convalescent SARS-CoV donor could react with one or more of the endemic HCoV spike proteins. However, little, if anything, is known about the functionalities and epitopes of these cross-reactive antibodies.

Coronavirus neutralizing antibodies target the trimeric spike (S) glycoproteins on the viral surface that mediate virus attachment and entry into the host cell. Several cryo-electron microscopy (cryo-EM) structures of trimeric spikes have been determined, with each comprising three S1 receptor binding subunits that collectively cap the trimeric S2 fusion subunit ^22–32^. Most antibodies neutralize coronavirus infection by binding to the receptor-binding subdomain of S1 and blocking receptor interactions. These antibodies are highly species specific and often strain specific due to a high sequence diversity in the receptor binding sites among coronaviruses, even for those that engage the same receptor (e.g. SARS-CoV and SARS-CoV-2) ^33^. Recently S1-targeting monoclonal antibodies have been identified that bind regions distal to the receptor binding site and that cross-neutralize coronaviruses within the *Sarbecovirus* subgenus ^21, 34–40^ with *in vivo* protective efficacy ^40, 41^, providing potential new leads for development of a pan-sarbecovirus vaccine. Neutralizing antibodies that broadly target coronaviruses from distinct (sub)genera are less likely to be found as the overall sequence identity among all coronaviruses spike proteins is low. Relative to the S1 subunit, the membrane-anchored S2 subunit, which mediates fusion of the viral and cellular membrane through receptor-induced conformational rearrangements, exhibits a higher level of protein sequence conservation across coronavirus spike proteins. Structural studies show exposed sites on the S2 base of the spike trimer that might be targeted by antibodies with cross-species specificity ^28, 42^.

Here we report first evidence of a class of S2-targeting antibodies with broad reactivity towards several human betacoronaviruses from three distinct subgenera, and characterized their antiviral activity, epitope and *in vivo* protective efficacy.

## Results

### Cross-reactivity of human monoclonal antibodies 28D9 and 1.6C7 to spike proteins of viruses in the *Betacoronavirus* genus

Five out of seven coronaviruses that are currently known to cause disease in humans belong to the *Betacoronavirus* genus, including SARS-CoV and SARS-CoV-2 (subgenus *Sarbecovirus*), MERS-CoV (subgenus *Merbecovirus*), and the endemic human coronaviruses HCoV-OC43 and HCoV-HKU1 (subgenus *Embecovirus*). To elicit and isolate betacoronavirus spike-targeting antibodies with cross-species specificity, we earlier immunized mice with trimeric spike ectodomains (S_ecto_) of three human infecting betacoronaviruses from different subgenera (HCoV-OC43, SARS-CoV and MERS-CoV) following a sequential immunization scheme as described in **Suppl. Fig. 1a**^34^. Transgenic H2L2 mice that encode the human immunoglobulin repertoire were used for immunization to develop fully human antibodies. We selected 203 hybridomas based on reactivity against at least one of the three spike antigens and screened their supernatants for cross-reactivity against five distinct betacoronavirus spike proteins by ELISA **(Suppl. Fig. 1b)**. Based on their broad ELISA reactivity profile towards multiple coronavirus spike proteins, we selected antibody 28D9 as well as a previously isolated MERS-S human monoclonal antibody 1.6C7 ^43^. Antibodies 1.6C7 and 28D9 were recombinantly expressed as human IgG1-isotype antibodies, and compared for their cross-reactivity profile, cross-neutralization capacity, mechanism of action, epitope characteristics and *in vivo* protective efficacy in subsequent experiments.

To determine the cross-reactivity of the 1.6C7 and 28D9 monoclonal antibodies we tested their binding to spike proteins of viruses in the *Betacoronavirus* genus by ELISA. Equimolar coating of the different antigens was verified by an antibody targeting the Strep-tag located at the C-termini of all antigens. The 1.6C7 and 28D9 antibodies reacted with spike ectodomains of betacoronaviruses from different subgenera including MERS-CoV (subgenus *Merbecovirus*), SARS-CoV and SARS-CoV-2 (subgenus *Sarbecovirus*), and HCoV-OC43 and MHV (subgenus *Embecovirus*), whereas no reactivity was seen with the spike ectodomain (S_ecto_) of HCoV-HKU1 (subgenus *Embecovirus*) **(Fig. 1a)**. Both mAbs bound equally well to MERS-S_ecto_ and OC43-S_ecto_, shown by the ELISA-based half maximal effective concentration (EC_50_) values. Compared to 1.6C7, 28D9 was superior in its cross-reactivity profile, with stronger binding to SARS-S_ecto_, SARS2-S_ecto_ and MHV-S_ecto_ **(Fig. 1a)**. Both antibodies reacted to S2 ectodomains (S2_ecto_) of MERS and MHV, indicating that their epitopes were located on the S2 fusion subunit of CoV spike proteins.

**Fig. 1.**
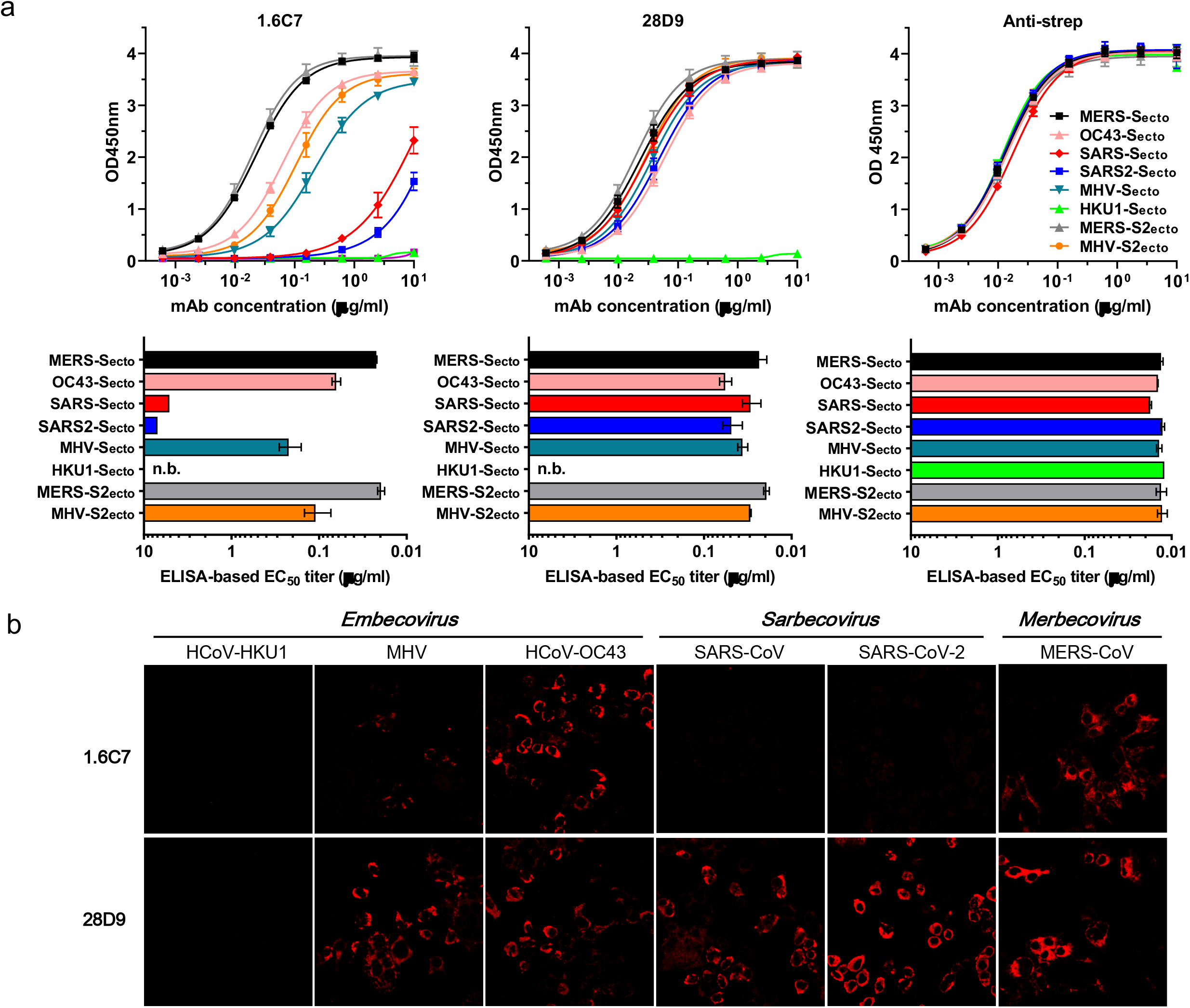
Cross-reactivity of human monoclonal antibodies 28D9 and 1.6C7 to spike proteins of viruses in the *Betacoronavirus* genus. **a.** ELISA binding curves (upper panels) and corresponding ELISA-based half-maximal effective concentrations (EC_50_) titers (lower panels) of mAbs 28D9 and 1.6C7 to Strep-tagged spike ectodomains (S_ecto_) and S2 ectodomains (S2_ecto_) of betacoronaviruses from different subgenera including MERS-CoV (subgenus *Merbecovirus*), SARS-CoV and SARS-CoV-2 (subgenus *Sarbecovirus)*, HCoV-OC43, HCoV-HKU1 and MHV (subgenus *Embecovirus*), coated at equimolar concentrations. Anti-strep mAb targeting the Strep-tagged antigens was used to corroborate equimolar plate coating. n.b., no binding. The average ± SD from two independent experiments with technical duplicates is shown. **b.** Binding of mAbs 1.6C7 and 28D9 to HEK-293T cells expressing GFP-tagged membrane-anchored full-length spike proteins of MERS-CoV, SARS-CoV, SARS-CoV-2, HCoV-OC43, HCoV-HKU1 and MHV detected by immunofluorescence assay. Cell nuclei in the overlay images were visualized by DAPI. The fluorescence images were recorded using a Leica SpeII confocal microscope.

Biolayer interferometry was used to characterize the binding kinetics and affinity of the cross-reactive antibodies to CoV S_ecto_ trimers immobilized on the surface of biosensors. The 1.6C7 mAb bound to the S_ecto_ of MERS-CoV, HCoV-OC43, SARS-CoV and SARS-CoV-2 with equilibrium dissociation constants (*K_D_*) of 0.58, 5.28 nM and 25.22 and 26.18 μM, respectively, whereas *K_D_*’s of the 28D9 mAb with these spike proteins were 0.72, 7.45, 3.17 and 5.96 nM, respectively **(Suppl. Fig. 2)**.

To assess whether the 1.6C7 and 28D9 antibodies could bind the full-length (i.c. membrane-anchored) version of spike proteins, we tested their reactivity to cells transiently expressing different (GFP-tagged) CoV spike proteins using immunofluoresence microscopy and flow cytometry. Binding of 1.6C7 was observed both in permeabilized and non-permeabilized cells expressing the spike proteins of HCoV-OC43, MERS-CoV and MHV, whereas 28D9 additionally bound to cells expressing the spike proteins of SARS-CoV and SARS-CoV-2 **(Fig. 1b, Suppl. Fig. 3 and Suppl. Table 1)**. No reactivity was seen for both antibodies to cells expressing the HCoV-HKU1 spike protein. These data demonstrate that both antibodies can react with full length spike proteins, with specificities that are consistent with the CoV S_ecto_ ELISA reactivities.

### Cross-neutralization capacity of mAbs 28D9 and 1.6C7 and mechanism of action

Next neutralizing activity by both antibodies against the targeted human coronaviruses was assessed. The 1.6C7 and 28D9 antibodies neutralize infection of MERS-S pseudotyped VSV (IC_50_ values of 0.39 and 0.13 μg/ml, respectively) as well as of authentic MERS-CoV (IC_50_: 0.083 and 0.93 μg/ml, respectively) **(Fig. 2a and b)**. No neutralization was seen against authentic SARS-CoV and SARS-CoV-2 by either of these antibodies, yet 28D9 displayed low levels of cross-neutralizing activity in the neutralization-sensitive pseudovirus system. 28D9 inhibited OC43-S, SARS-S and SARS2-S pseudotyped VSV infection with IC_50_ values of 68.4, 60.5 and 45.3 μg/ml, respectively **(Fig. 2a)**.

**Fig. 2.**
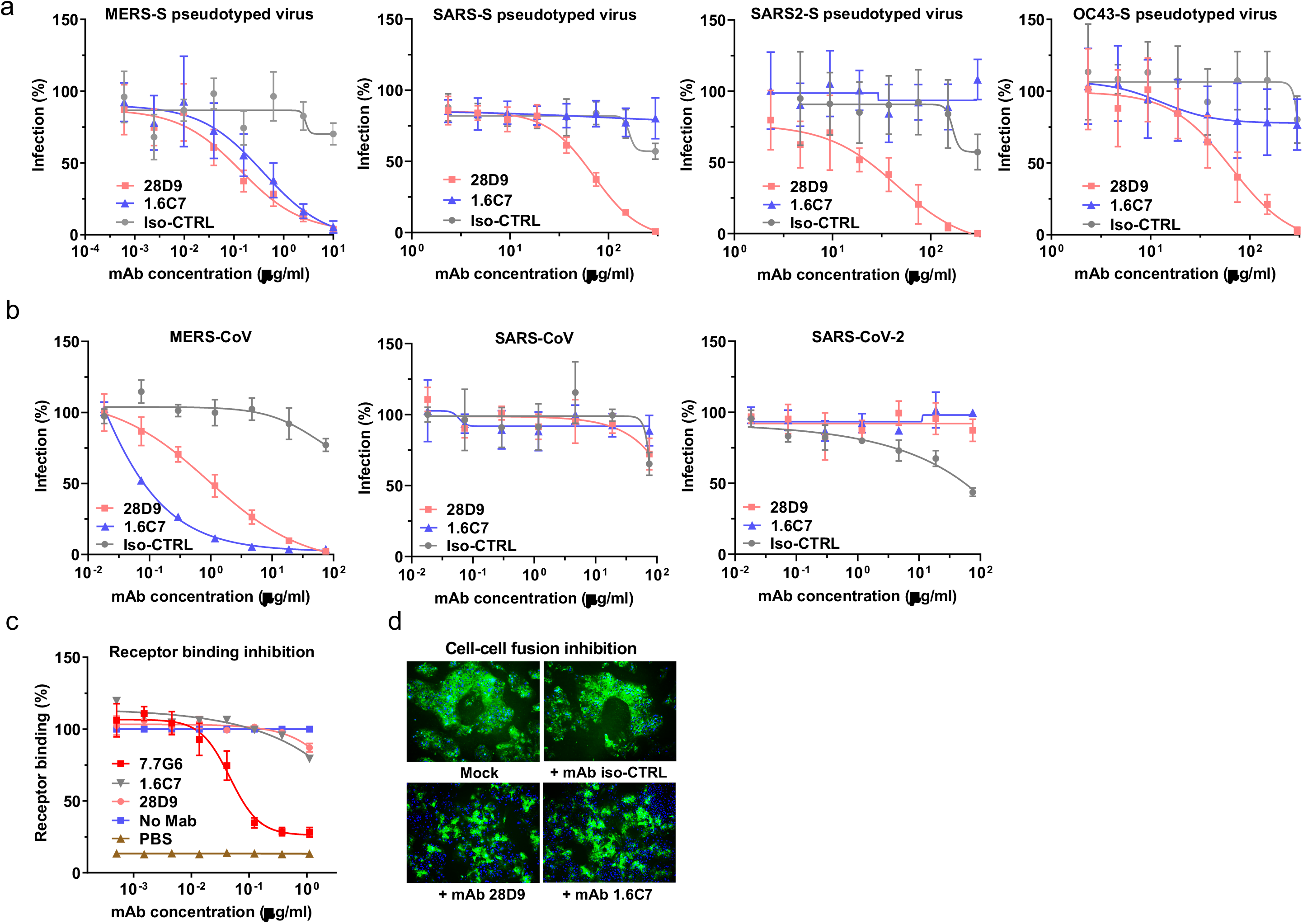
Cross-neutralization capacity of mAbs 28D9 and 1.6C7 and mechanism of action. **a.** Antibody-mediated neutralization of infection of luciferase-encoding VSV particles pseudotyped with spike proteins of MERS-CoV, SARS-CoV, SARS-CoV-2 and HCoV-OC43. Pseudotyped VSV particles pre-incubated with antibodies at indicated concentrations were used to infect VeroCCL81 cells (MERS-S pseudotyped VSV), VeroE6 cells (SARS-S and SARS2-S pseudotyped VSV) or HRT-18 cells (OC43-S pseudotyped VSV) and luciferase activities in cell lysates were determined at 20 h post transduction to calculate infection (%) relative to non-antibody-treated controls. The average ± SD (n = 3) from at least two independent experiments performed is shown. Iso-CTRL: an anti-Strep-tag human monoclonal antibody was used as an antibody isotype control. **b.** Antibody-mediated neutralization of MERS-CoV, SARS-CoV and SARS-CoV-2 infection. Neutralization of authentic viruses was performed using a plaque reduction neutralization test (PRNT) on VeroCCL81 cells (MERS-CoV) or VeroE6 (SARS-CoV and SARS-CoV-2) as described earlier ^65, 66^. The experiment was performed with triplicate samples, the average ± SD is shown. **c.** ELISA-based receptor binding inhibition assay. MERS-S_ecto_ pre-incubated with serially diluted mAbs was added to ELISA plates coated with soluble human DPP4. Binding of MERS-S_ecto_ to DPP4 was detected using an HRP-conjugated antibody recognizing the C-terminal Strep-tag on MERS-S_ecto_. The average ± SD from two independent experiments with technical duplicates is shown. **d.** Cell-cell fusion inhibition assay. Huh-7 cells - transfected with plasmid expressing (GFP-tagged) MERS-CoV S were pre-incubated in the presence or absence of 1.6C7 and 28D9, or an irrelevant iso-type control antibody (Iso-CTRL), and then treated with trypsin to activate the membrane fusion function of the MERS-CoV S protein. Formation of MERS-S mediated syncytia was visualized by fluorescence microscopy. Merged images of MERS-S-GFP expressing cells (green) and DAPI-stained cell nuclei (blue) are shown. The experiment was performed twice, data from a representative experiment is shown.

To understand the mechanism of virus neutralization, we assessed antibody interference with spike-mediated receptor binding and membrane fusion activity. In line with our earlier observations for 1.6C7 ^43^, we found that 28D9 inhibits MERS-S driven cell-cell fusion but does not impede MERS-S_ecto_/DPP4 receptor interaction **(Fig. 2c and d)**, suggesting that both S2-targeting antibodies prevent the membrane fusion function of the spike S2 subunit that is required for infection.

### mAbs 1.6C7 and 28D9 target a linear epitope located in the stem region of the S2 fusion subunit

Both antibodies were tested for competitive binding to MERS-S_ecto_ using biolayer interferometry. MERS-S_ecto_ binding by 1.6C7 antibody compelety prevented binding of 28D9, and vice versa, suggesting that the antibodies target overlapping epitopes **(Fig. 3a)**. To assess the epitope conformational specificity, we compared the ELISA reactivity of 1.6C7 and 28D9 to untreated MERS-CoV S_ecto_ antigen versus antigen that was heat-denatured in the presence of SDS and DTT. Both antibodies reacted equally well with non-denatured and denatured proteins, unlike a MERS-S1 targeting antibody 7.7G6 ^43^ that only reacted with non-denatured antigen. These data indicate that 1.6C7 and 28D9 likely bind a linear, contiguous sequence of amino acids in MERS-S **(Fig. 3b)**.

**Fig. 3.**
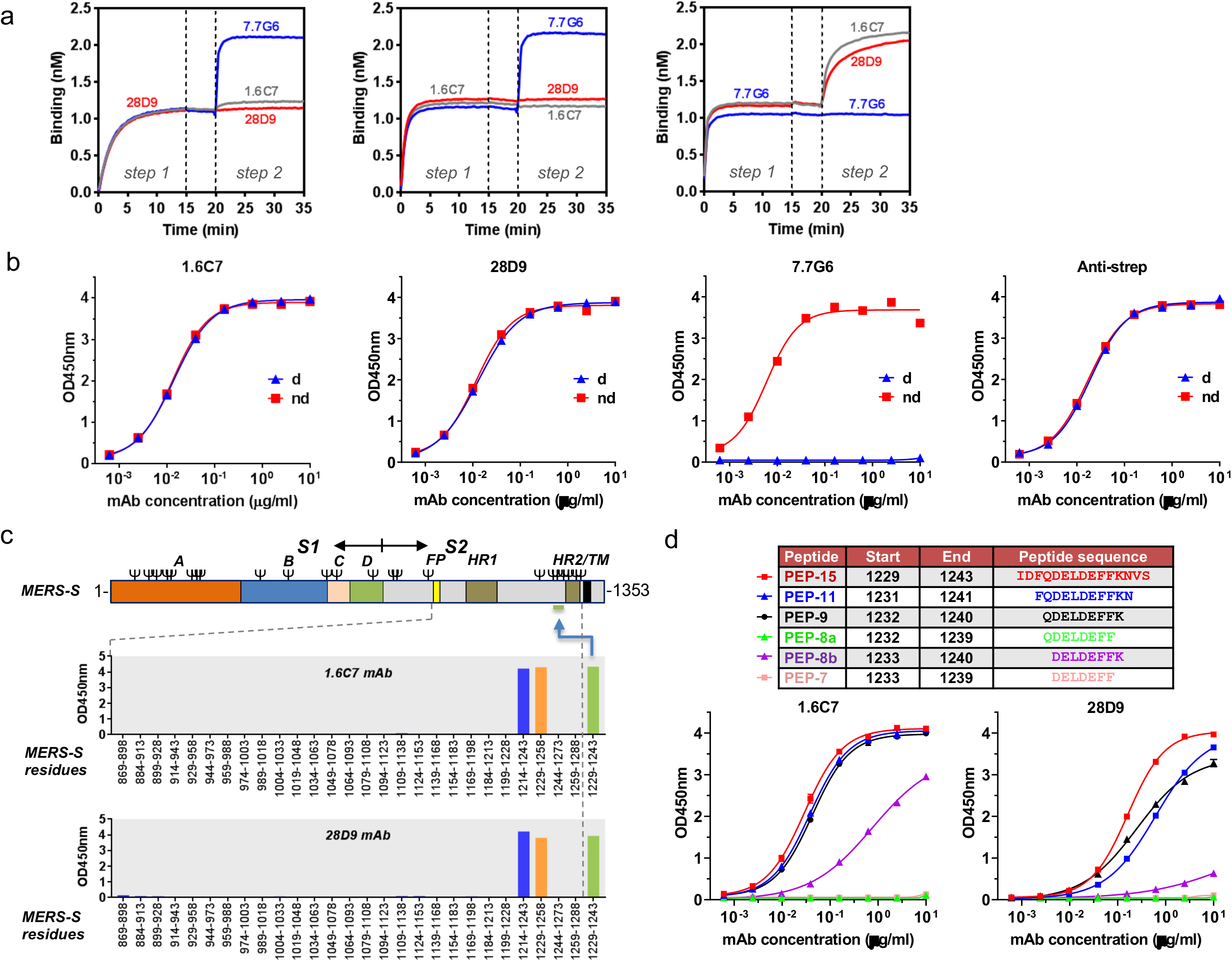
mAbs 1.6C7 and 28D9 target a linear epitope located in the stem region of S2 fusion subunit. **a.** Antibody binding competition analysed by biolayer interferometry. Immobilized MERS-S_ecto_ antigen was saturated in binding with a given mAb (step 1) and then exposed to binding by a second mAb (step 2). Additional binding of the second antibody indicates the presence of an unoccupied epitope, whereas lack of binding indicates epitope blocking by mAb1. As a control, the first mAb was also included in the second step to check for self-competition. Competitive binding was tested for the S2-targeting 1.6C7 and 28D9 antibodies and a MERS-S1 antibody control (7.7G6) ^43^. **b.** 1.6C7 and 28D9 recognize a linear epitope. ELISA binding curves of 1.6C7 and 28D9 to untreated MERS-S_ecto_ (non-denatured: ‘nd’) versus MERS-S_ecto_ that was heat-denatured in the presence of SDS and DTT (denatured: ‘d’). Two antibodies targeting the MERS-S1 domain (7.7G6) and the 8-residue long linear Strep-tag epitope (anti-strep) were used as controls. **c**. 1.6C7/28D9 epitope maps to a 15-aa long stem region upstream of HR2 in MERS-S. ELISA-reactivity of 1.6C7 and 28D9 to a peptide library of 30-amino acid long peptides (with 15-a.a. overlap) covering the conserved C-terminal part of the MERS-S_ecto_ (residues 869-1,288). Both antibodies reacted with two peptides (blue and orange bars), and with a peptide corresponding to their 15-a.a. long overlap (green bar; MERS-S residues 1,229-1,245). The position of the epitope containing region is indicated in the MERS-S protein schematic with the spike subunits (S1 and S2), S1 domains (A through D), fusion peptide (FP), heptad repeat 1 (HR1), heptad repeat 2 (HR2) and transmembrane domain (TM) annotated. **d**. The 1.6C7/28D9 epitope maps to a 8-aa long peptide ‘DELDEFFK’ detected by ELISA. ELISA binding curves of 1.6C7 and 28D9 to N- and C-terminally truncated versions of the 15-mer peptide fragment of MERS-S. Data from a representative experiment with technical duplicates are shown.

As both antibodies may bind a linear epitope, we aimed to map their epitope location by ELISA using an array of overlapping synthetic peptides (30-mer peptides with an overlap of 15 residues) covering the conserved C-terminal part of the MERS-S2_ecto_ (residues 869-1,288). Both antibodies appeared to bind a 15-residue-long region (MERS-S residues 1,229-1,243) positioned just upstream of the heptad repeat 2 (HR2) region in the MERS-S2 fusion subunit **(Fig. 3c)**. Analysis of 1.6C7/28D9 antibody binding to N- and C-terminally truncated versions of the 15-mer peptide fragment designated the D^^1233^^ELDEFFK^^1240^^ MERS-S protein segment as the minimal epitope region still affording some binding by both antibodies **(Fig. 3d)**.

### Fine mapping the 1.6C7 and 28D9 antibody binding sites on the spike protein by mutagenesis

To map residues critical for binding by the 1.6C7/28D9 antibodies, we performed ELISA-based epitope alanine scanning mutagenesis on the 15-mer spike peptide fragment comprising the linear epitope. The alanine scanning analysis of the 15-mer peptide fragment defined three residues (D^^1236^^, F^^1238^^ and F^^1239^^) as critical to binding by both antibodies **(Fig. 4a and Suppl. Fig. 4)**, with another four residues (D^^1233^^, E^^1234^^, L^^1235^^ and E^^1237^^) that contribute to binding by 28D9 and - to a lesser extent - by 1.6C7. To corroborate the epitope alanine scanning data, we assessed ELISA reactivity of the 1.6C7/28D9 antibodies to single site MERS-S_ecto_ mutants containing alanine substitutions in the core epitope region. Consistent to the peptide alanine scanning, alanine substitution of the three residues D^^1236^^, F^^1238^^ and F^^1239^^ abrogated MERS-S_ecto_ binding by both antibodies, but not that of the anti-MERS-S1 control antibody 7.7G6 **(Fig. 4c)**.

**Fig. 4.**
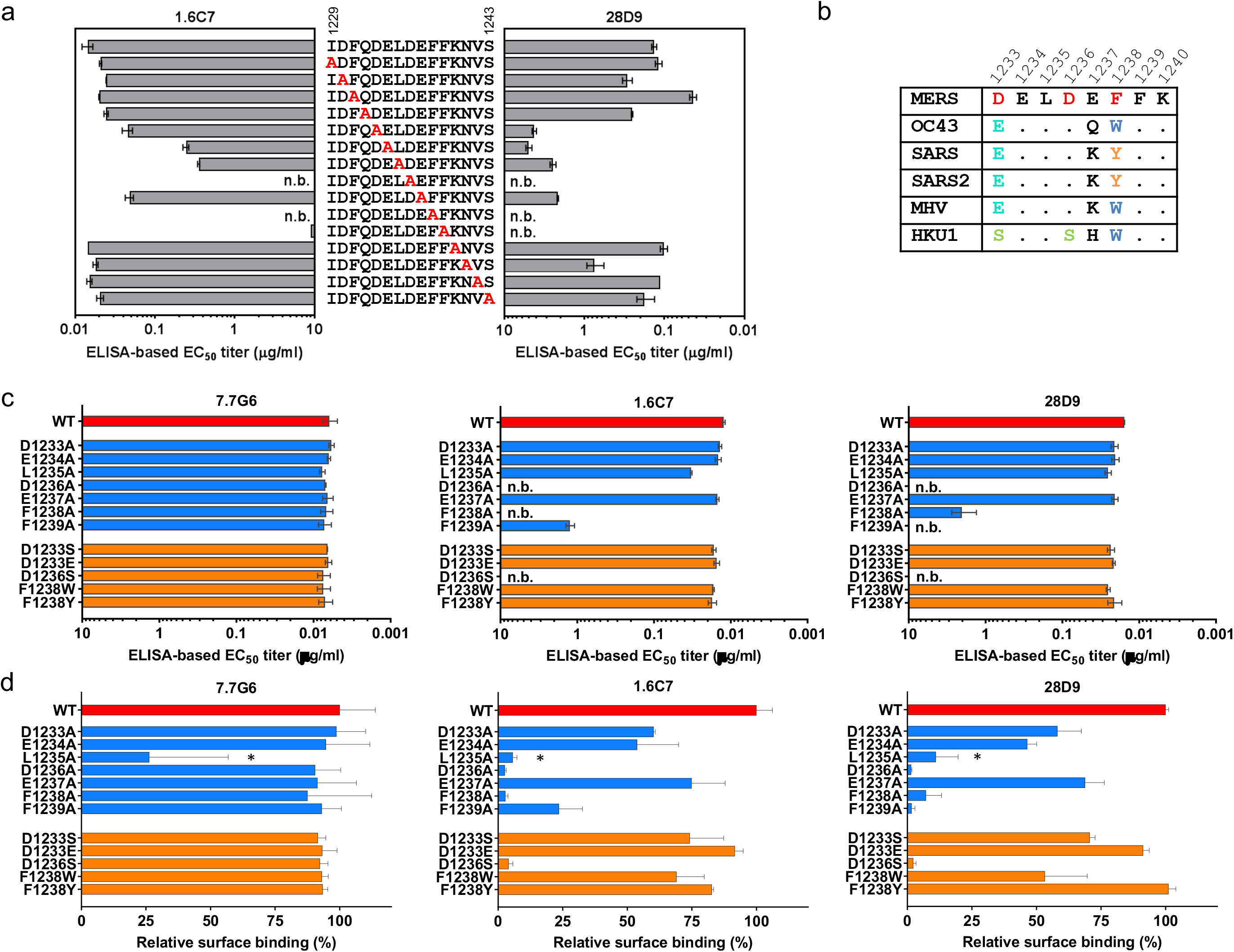
Fine mapping the 1.6C7 and 28D9 antibody binding sites on the spike protein by mutagenesis. **a.** ELISA-based epitope alanine mutagenesis on the 15-mer spike peptide fragment comprising the linear 1.6C7 and 28D9 epitope, shown by half-maximum effective concentration (EC_50_) titers (μg/ml). The average ± SD from two independent experiments performed is shown. **b.** Sequence alignment of 1.6C7/28D9 epitope region of MERS-CoV, HCoV-OC43, SARS-CoV, SARS-CoV-2, MHV and HCoV-HKU1. **c.** Spike protein ectodomain single site mutagenesis to delineate 1.6C7 and 28D9 antibody binding sites. ELISA-based EC_50_ titers (μg/ml) of 1.6C7/28D9 binding to MERS-S_ecto_ mutants containing single amino acid substitutions in the core epitope region are indicated on the left. Anti-MERS-S1 control antibody 7.7G6 was used to control the expression level of all mutants. n.b., no binding. The average ± SD from at least two independent experiments performed is shown. **d**. Binding of 28D9 and 1.6C7 antibodies to cell surface expressing (GFP-tagged) MERS-S mutants containing single amino acid substitutions in the core epitope region detected by flow cytometry. Antibody binding was detected using AlexaFluor 594 conjugated secondary antibody. Relative surface binding was determined by calculating the percentage of GFP^+^/Alexa Fluor 594^+^ cells over GFP^+^ cells. Anti-MERS S1 antibody 7.7G6 was used to control the cell surface expression levels of all single-site mutants. The asterisk indicates reduced cell surface expression of the L1235A mutant. n.b., no binding. The average ± SD from two independent experiments performed is shown.

To assess the individual contribution to antibody binding of identified residues in the context of the membrane-anchored full-length spike protein, a flow cytometry-based assay was performed measuring antibody binding to cell-surface expressed MERS-S mutants **(Fig. 4d)**. The anti-MERS-S1 7.7G6 antibody was used to control cell surface expression levels of the single-site mutants. With the exception of the L^^1235^^A mutant, all mutant spike proteins displayed surface expression levels similar to wildtype. Alanine substitutions of residues in the MERS-S core epitope region (D^^1233^^ELDEFFK^^1240^^) reduced binding by both antibodies to a varying extent, although analysis of binding data for the L^^1235^^A mutant was compromised by reduced cell surface expression level. Consistent with the ELISA-based analysis **(Fig. 4a and c)**, mutation of D^^1236^^, F^^1238^^ and F^^1239^^ most strongly affected binding by both antibodies. Both antibodies showed similar binding patterns in all three antigen binding assays **(Fig. 4a, c-d and Suppl. Fig. 4)**, yet subtle differences in the reactivities were observed that may rationalize differences between these two antibodies and their cross-spike binding properties.

The identified residues in the spike epitope region (D^^1233^^ELDEFFK^^1240^^ in MERS-S) contributing to 1.6C7/28D9 binding are not fully conserved in the targeted betacoronavirus spike proteins, with variations found in three residues (D^^1233^^, D^^1236^^ and F^^1238^^ in MERS-S) that are key to binding by both cross-reactive antibodies **(Fig. 4b)**. To assess contribution of these amino acid variations to antibody reactivity, single site substitutions were made in MERS-S. Soluble and full-length D^^1233^^S, D^^1233^^E, D^^1236^^S, F^^1238^^W and F^^1238^^Y MERS-S mutants were tested for antibody binding by respectively ELISA **(Fig. 4c)** and flow cytometry **(Fig. 4d)**, as described above. Substitution of D^^1233^^ in MERS-S to serine (S, found in HKU1-S) or glutamic acid (E, found in SARS-, SARS2-, OC43- and MHV-S) still allowed efficient binding by both antibodies. In addition, but contrary to the MERS-S F^^1238^^A mutant, the MERS-S mutants in which F^^1238^^ was replaced by a tryptophan (W, found in HKU1-, OC43- and MHV-S) or tyrosine (Y, found in SARS- and SARS2-S) maintained antibody binding reactivity. These data indicate a degree of sequence variability that is allowed in the epitope region without compromising antibody binding. Particularly, different aromatic residues (Phe, Trp and Tyr) seem to be permitted at residue position 1238. Conversely, the D^^1236^^S substitution in MERS-S (found in HKU1-S) fully abrogated binding by both antibodies. The reciprocal S^^1236^^D substitution in HKU1-S was sufficient to rescue binding by 28D9, and to a lesser extend by 1.6C7 **(Suppl. Fig. 5)**. These data rationalize the ability of both antibodies to bind spike proteins of MERS-CoV, SARS-CoV, SARS-CoV-2, HCoV-OC43 and MHV as well as the inability of both antibodies to bind the HCoV-HKU1 spike protein. Evidently, epitope mapping of the two antibodies raised in two different animal immunization experiment revealed that they target the same epitope. We identified a third cross-reactive monoclonal antibody from an independent mouse immunization experiment (DNA+protein immunization), that targets the same site on betacoronavirus spike proteins **(mAb 18H2, Suppl. Fig. 6)**, indicating that this epitope is immunogenic and efficient in the induction of cross-reactive antibodies.

Sequence analysis of the variable regions of the 1.6C7, 28D9 and 18H2 antibodies identified that their heavy and light chains were derived from the IGHV6-1 and IGKV4-1 germline precursor, respectively. Six somatic hypermutation were found in the VH region for 28D9, and five for 1.6C7. Seven somatic hypermutation were found in the VL region for 28D9, and two for 1.6C7 **(Suppl. Fig. 7)**.

Glycans on antigens are frequently found to be a determinant for antibody binding. We found that antigen deglycosylation resulted in lower binding of 28D9 to OC43-S_ecto_, MHV-S_ecto_ and SARS-S_ecto_ using western blotting **(Supl. Fig. 8a)**, in contrast to 1.6C7. We tested the involvement of an N-linked glycan located one residue downstream of the core epitope region (D^1233^ELDEFFK^1240^) in MERS-S. This N-glycosylation sequon (NxS/T) is conserved among betacoronavirus spike protein orthologous **(Fig. 5a)**. Deletion of the N^1241^ glycosylation site in MERS-S_ecto_ did not impair binding by both antibodies in ELISA **(Suppl. Fig. 8b)**. In contrast, deletion of the orthologous glycosylation site (N^1241^) in OC43-S_ecto_ fully abrogated ELISA-reactivity of 28D9, whereas that of 1.6C7 was virtually unchanged. These data collectively indicate that epitope features required for 28D9 binding varied among the betacoronavirus spike proteins, with differential involvement of an N-linked glycan in antigen binding.

**Fig. 5.**
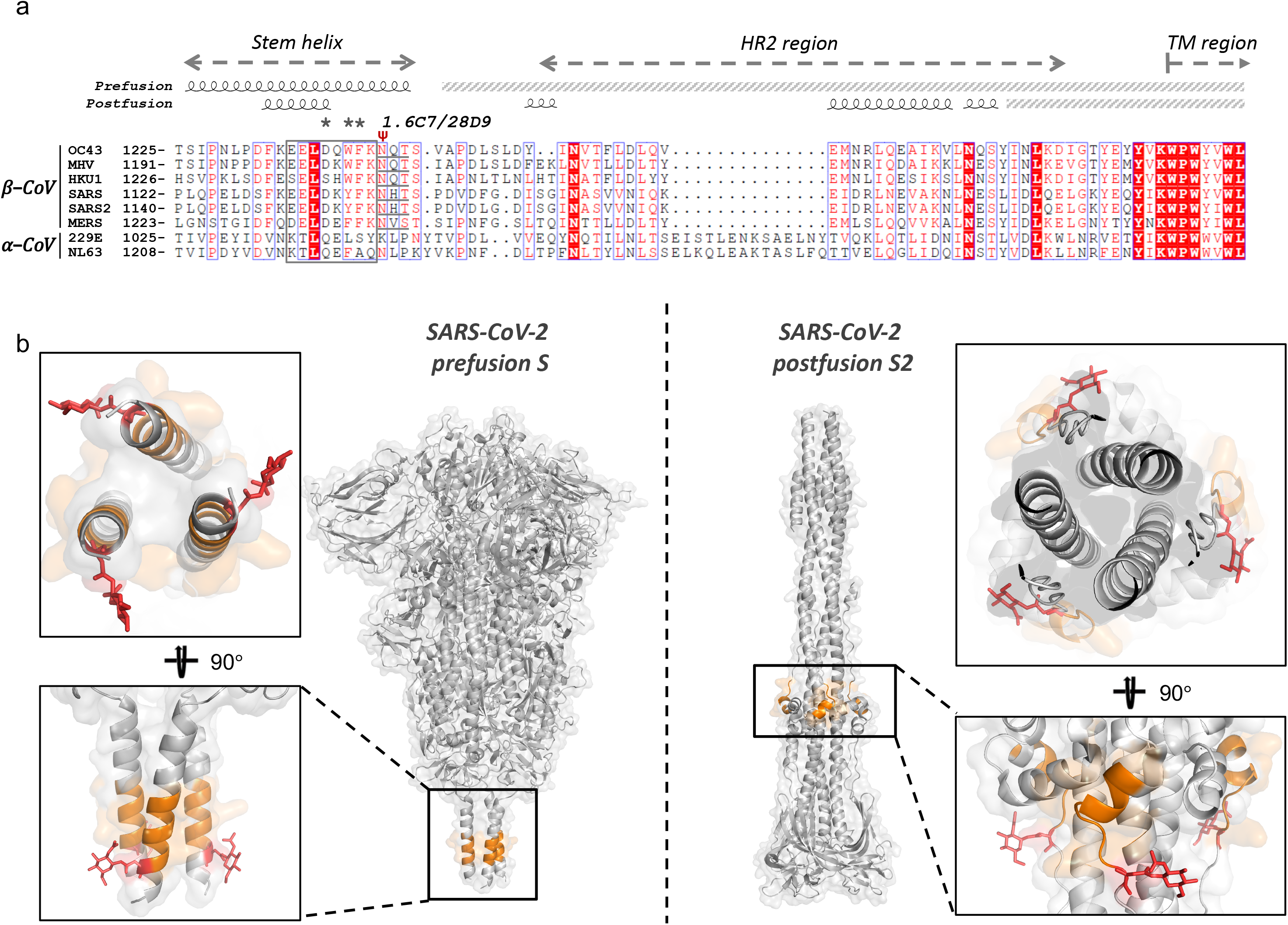
1.6C7 and 28D9 bind the membrane proximal stem helix of the coronavirus spike protein. **a.** Sequence alignment of spike protein region of alpha- and betacoronaviruses encompassing the 1.6C7/28D9 epitope region. The 28D9 and 1.6C7 core epitope region is outlined by a rectangle box and residues critical for antibody binding are annotated by asterisks. A conserved glycosylation sequon (NxS/T) found in betacoronavirus spike proteins - one amino acid downstream of the core epitope - is underlined and annotated (ψ). The stem helix, heptad repeat region 2 (HR2) and the start of the transmembrane domain (TM) are indicated. Secondary structural elements of the SARS-CoV prefusion spike (PDB: 6XR8) and postfusion S2 (PDB: 6XRA) structures are visualized using ESPript 3.0 (http://espript.ibcp.fr/ESPript/ESPript/). **b**. Structures of the SARS-CoV-2 spike (PBD: 6XR8) and S2 (PDB: 6XRA) in pre- and postfusion conformation, respectively. Structures are indicated as a grey cartoon with transparent surface presentation, and the segment corresponding to the stem helix epitope colored in orange. Insets: zoom-in sections of the epitope region in both structures in two different orientations with the conserved *N*-glycan highlighted in red.

### 1.6C7 and 28D9 target the membrane proximal stem helix of the coronavirus spike protein

Although the 28D9/1.6C7 epitope region was present in five betacoronavirus spike protein ectodomains of which cryo-EM prefusion structures were elucidated, the C-terminally located epitope region was not or only poorly resolved in the cryo-EM maps, indicative for conformational flexibility of the epitope region ^22, 32, 44^. However, the epitope was recently revealed in trimeric pre- and postfusion SARS-CoV-2 spike structures that were reconstructed by cryo-EM using purified full-length spike protein ^31^. In the prefusion SARS-CoV-2 spike, the epitope region (E^^1150^^ELDKYFK^^1157^^ in SARS2-S) is part of a 20-residue-long α-helix at the membrane-proximal base of the molecule **(Fig. 5b)**. The corresponding region in HCoV-NL63 spike protein was designated as ‘stem’ helix ^44^. The stem helices of three S protomers align along the three-fold symmetry axis and form a helical bundle accessible for antibody binding, and which displays a relatively high level of conserved surface exposed residues across betacoronavirus spike proteins **(Suppl. Fig. 9)**. Remarkably, both antibodies can also bind the postfusion spike structure as demonstrated by their ability to bind the MHV S2_ecto_ **(Fig. 1a)** that was earlier solved in its postfusion structure ^45^. In the S2 postfusion structure, the N- and C-terminal residues of the stem helix are uncoiled whereas the middle part (F^^1148^^KEELD^^1153^^ in SARS2-S) is still folded as a short, surface exposed helix, which docks perpendicular to the central coiled coil ^31, 45^.

### Antibodies towards the stem helix epitope are elicited during natural infection

We next analysed whether antibodies towards the stem helix epitope or other linear epitopes in the spike protein are elicited during natural MERS-CoV infection in humans and dromedary camels. Spike protein peptide microarray analysis with five human and four dromedary camel MERS-CoV-positive sera on 905 overlapping peptides covering the entire spike protein ectodomain (MERS-S residues 1-1,296) identified sixteen linear core epitopes **(Fig. 6 and Suppl. Fig. 10)**. Five of them were concentrated in the ±80-residue long region upstream of the spike transmembrane domain comprising the HR2 region, which undergoes extensive structural rearrangements during fusion. One of these five peptides (D^^1230^^FQ<u>DELDEFFK</u>NVS^^1243^^) is overlapping the epitope core region (underlined in sequence) of the 1.6C7/28D9 antibodies **(Fig. 6c)** and is – compared to other identified linear peptide epitopes - recognized most frequently by MERS-positive sera from humans (3/5) and dromedary camels (2/4) **(Fig. 6c and Suppl. Fig. 10)**. These data indicate that antibodies towards this epitope are efficiently elicited not only in spike-immunized mice but also during natural infection of humans and dromedary camels.

**Fig. 6.**
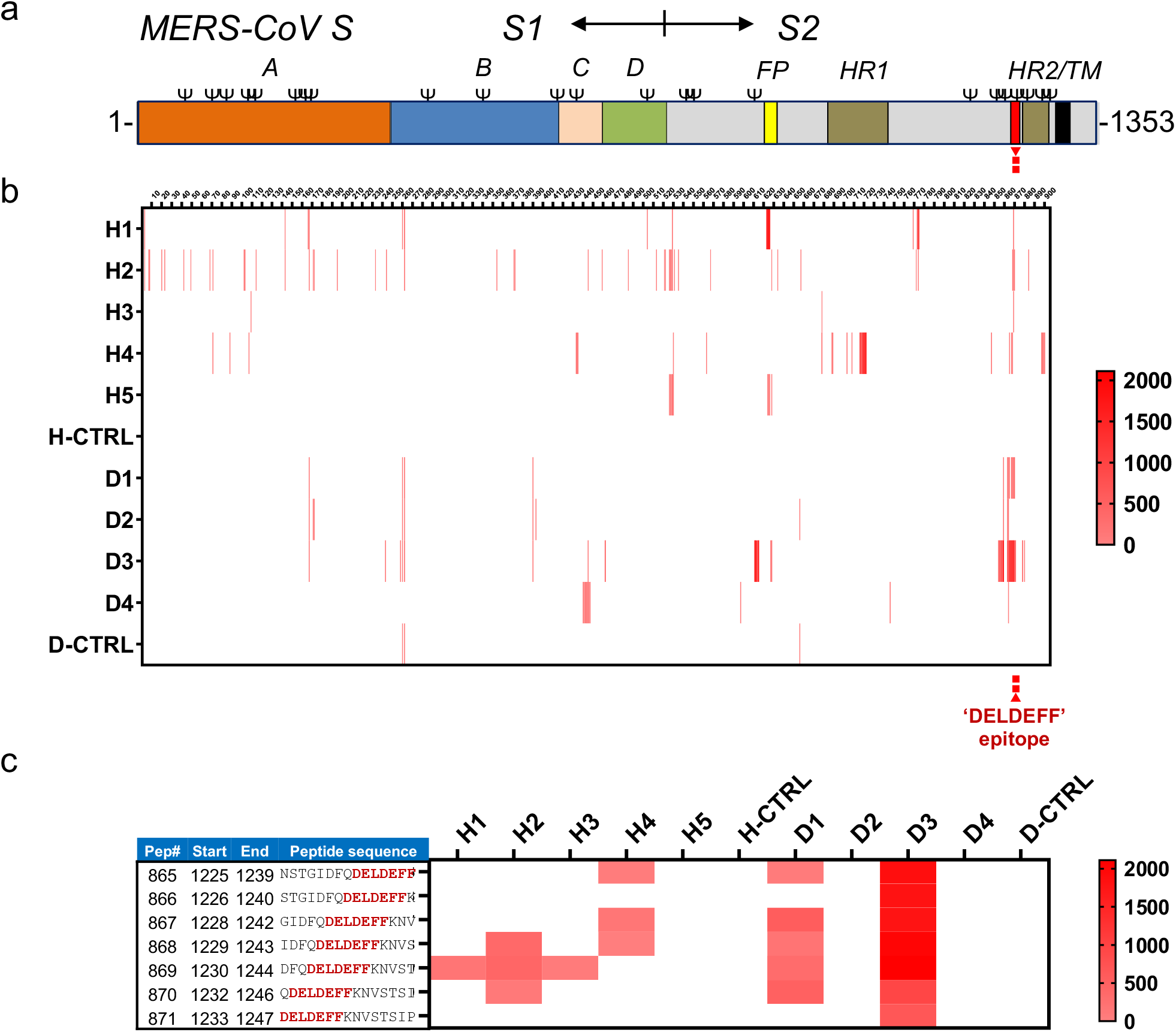
Antibodies towards the stem helix epitope are elicited during natural infection. **a.** Schematic representation of the MERS-CoV spike protein. The spike subunits (S1 and S2), S1 domains (A through D), fusion peptide (FP), heptad repeat 1 (HR1), heptad repeat 2 (HR2) and transmembrane domain (TM) are annotated. **b.** Spike protein peptide microarray analysis using MERS-positive human and dromedary camel sera. 905 overlapping peptides covering the entire MERS-CoV S ectodomain (residues 1-1,296) were synthesized with an offset of one or two residues. The binding of five convalescent MERS-positive human (H1 to H5) and four dromedary camel (D1 to D4) sera to the peptide library, as well as a MERS-negative serum from human (H-CTRL) or camel (D-CTRL) was assessed in a PEPSCAN-based ELISA (Lelystad, The Netherlands). Cumulative heatmap of signal intensities for individual peptides are shown. Signal intensities increase from light reddish to red, whereas white corresponds to background signal. **c.** Reactivity of the human and dromedary sera to peptides covering the 1.6C7/28D9 epitope region (epitope core sequence highlighted in red).

### Antibody mediated protection of mice challenged with MERS-CoV or SARS-CoV

To assess the *in vivo* protection capacity of antibodies targeting the stem-helix epitope, we tested the prophylactic and therapeutic activity of the 1.6C7 mAb against lethal MERS-CoV challenge using the K18-*hDPP4* transgenic mouse model expressing human DPP4 ^46^. 20-30-week-old mice were injected with 50 μg of antibody (equivalent to 1.8 mg mAb per kg body weight) by intraperitoneal injection 24 hours before (pre-) or 24 hours after infection (post-) with a lethal dose of MERS-CoV. A highly potent MERS-CoV neutralizing antibody 7.7G6, and an irrelevant IgG1 isotype control antibody ^43^ were taken along as controls **(Fig. 7a)**. The percentage of survival and weight change was monitored daily for 10 days. In isotype control pre- and post-treated mice, MERS-CoV causes lethal disease with 100% lethality between 8 and 10 days and showed significant weight loss. In contrast, both pre- and post-treatment of 1.6C7 protected mice from death, and that of 7.7G6 protect 80-100% of mice from death. Relative to isotype control treated mice, mice treated with either 7.7G6 or 1.6C7 showed reduced weight loss **(Fig. 7a)**.

**Fig. 7.**
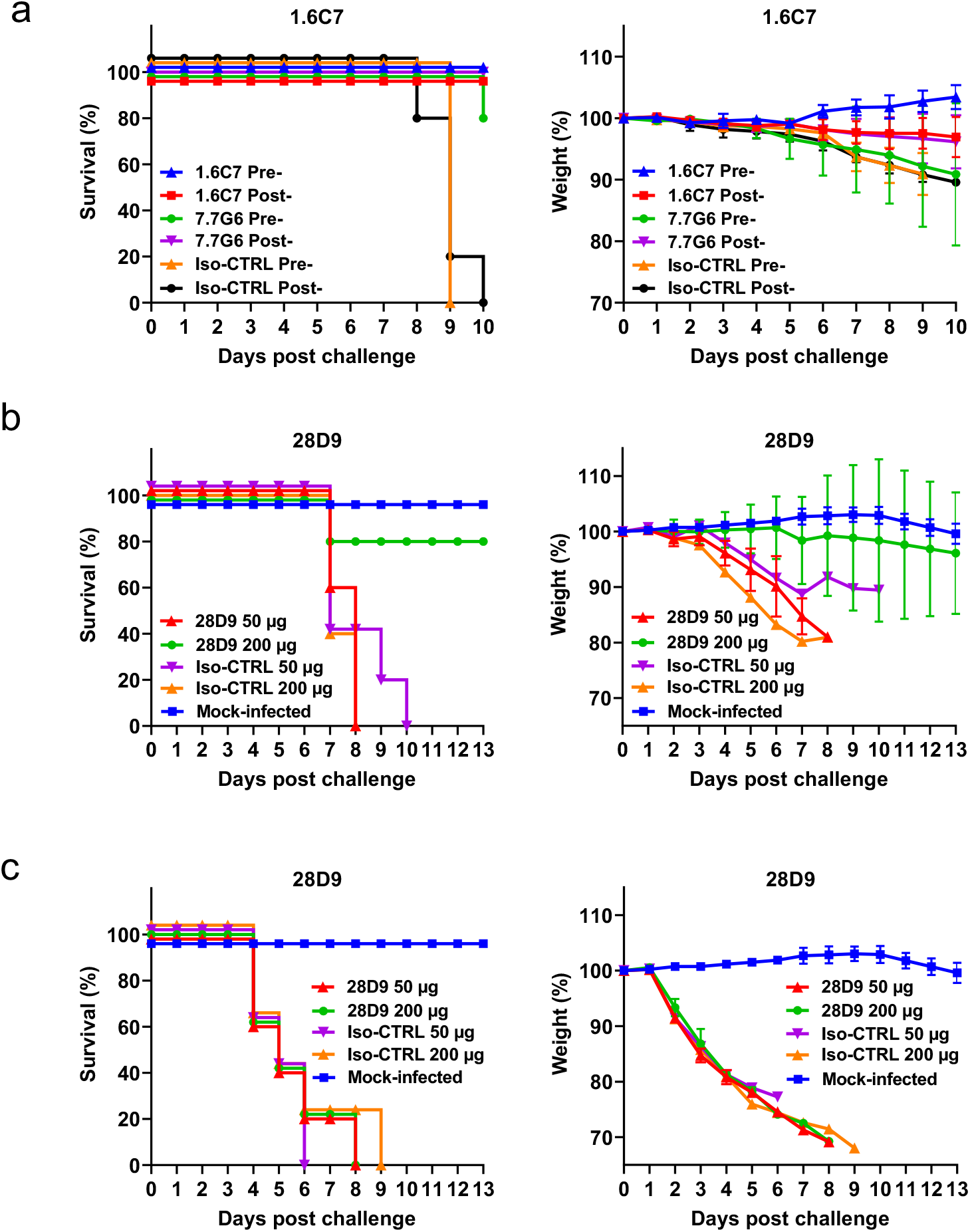
Antibody mediated protection of mice against lethal MERS-CoV/SARS-CoV challenge. **a.** The *in vivo* prophylactic and therapeutic activity of the 1.6C7 mAb against lethal dose MERS-CoV challenge was tested in the K18 transgenic mouse model expressing human DPP4 ^46^. A potent neutralizing MERS-S1 antibody (7.7G6) or an irrelevant IgG1 control antibody was taken along. Eight 20-30-week-old mice were injected with 50 μg of antibody (equivalent to 1.8 mg mAb/kg body weight) by intraperitoneal injection 24 hours before (pre-) or 24 hours after (post-) intranasal infection with a lethal dose of MERS-CoV. Survival rates (left) and weight loss (right, expressed as a percentage of the initial weight) were monitored daily until 10 days post-inoculation. **b.** Prophylactic efficacy of 28D9 against MERS-CoV infection. Five 20-week-old K18 mice were mock-infected or injected with 50 or 200 μg of 28D9 (equivalent to 1.8/7.2 mg mAb/kg body weight) or isotype control antibody by intraperitoneal injection 24 hours before intranasal infection with a lethal dose of MERS-CoV. **c.** Prophylactic efficacy of 28D9 against SARS-CoV infection. Five 16-week old Balb-C mice were mock-infected or administered with the 50 or 200 μg of 28D9 or isotype control antibody via intraperitoneal injection 24 hours before intranasal infection with a lethal dose of mouse adapted SARS-CoV. Survival rates and weight loss (expressed as a percentage of the initial weight) were monitored daily until 13 days post-inoculation.

To investigate the reduction in pathology and viral loads, lungs of mice were harvested on day 3 (3 animals) and 8-10 (5 animals) post infection. In isotype control treated mice, no obvious difference of viral RNA was observed compared to non-antibody treated (Mock-treated) mice. However, mice treated with 1.6C7 before or after virus challenge, demonstrated 1-2 log reduction in viral RNA titers on day 3 post exposure, whereas a 2-3 log reduction was seen at 8-10 dpi **(Suppl. Fig. 11a)**. A similar reduction was seen for the isolation of infectious virus at day 3, whereas at day 8-10 post exposure no infectious virus could be isolated from the lungs of the 1.6C7 pre- and post-exposure treated mice **(Suppl. Fig. 11a)**. Mice treated with 1.6C7 before and after infection showed greatly reduced lung pathology and inflammation following MERS-CoV infection at day 3 and 8-10, consistent with viral RNA levels and virus titers determined for these mice **(Suppl. Fig. 11b)**.

We next tested the *in vivo* protection activity of the 28D9 mAb against MERS-CoV and SARS-CoV challenge. Mice (n=5) were injected with two doses (50 or 200 μg) of 28D9 or an isotype control antibody by intraperitoneal injection 24 hours before intranasal infection with a lethal dose of MERS-CoV or of SARS-CoV **(Fig. 7b and c),** respectively. The percentage of survival and weight change of mice was monitored daily for 13 days. Whereas no protection was seen against SARS-CoV challenge, four out of five animals infused with 200 μg 28D9 antibody survived MERS-CoV infection and did not display significant weight loss.

## Discussion

We report two human monoclonal antibodies that are able to bind the spike proteins of betacoronaviruses from three different subgenera. Both antibodies were found to target a conserved and immunogenic epitope in membrane-proximal stem region on the S2 fusion subunit, that is also recognized by the immune system during natural infection. Antibodies targeting this antigenic site prevented mortality and disease in mice upon MERS-CoV infection, demonstrating the relevance of the epitope for providing protection.

The observed antibody cross-reactivities are remarkable since the spike protein sequences of the five targeted betacoronaviruses (MHV, HCoV-OC43, MERS-CoV, SARS-CoV and SARS-CoV-2) are highly divergent and share only about 15% overall protein sequence identity with 26% identity across the more conserved S2 fusion subunit. Three residues in the stem helix epitope region appeared to be critical for antibody binding. Two of those appeared fully conserved in the targeted betacoronavirus spike sequences (D^^1236^^ and F^^1239^^ in MERS-S), whereas only aromatic amino acids residues were found at the third position (F^^1238^^ in MERS-S), indicative of functional conservation. Location of the epitope was corroborated by the gain-of-function binding for the HCoV-HKU1 spike protein upon serine substitution at MERS-S equivalent position D^^1236^^. The small number of (functionally) conserved key residues targeted by both antibodies and the involvement of a conserved N-linked glycan may rationalize their binding to these highly divergent antigens ^47, 48^.

We showed that the stem-helix targeting antibodies effectively blocked MERS-CoV infection in cultured cells and provided prophylactic and therapeutic protection to mice against a lethal dose MERS-CoV challenge. This epitope was hitherto not described as a target of neutralizing antibodies, although monoclonal antibodies with neutralizing activity were found to target the sequences flanking the stem-helix epitope, including the downstream HR2 region ^49–51^. Membrane fusion by the S2 subunit is mediated through extensive conformational rearrangements, particularly in the epitope containing region. Elucidation of pre- and post-fusion spike structures reveals that dissociation of the three epitope-containing stem-helices in the prefusion spike trimer is required for formation of the highly stable postfusion structure ^31, 45, 52, 53^. Though the precise mechanism of neutralization by these two antibodies remains to be defined, at least two mechanisms of neutralization are conceivable: the binding of antibodies may destabilize the bundle of stem helices in the prefusion spike protein initiating premature spike activation. Alternatively, antibody binding may obstruct six-helix bundle formation of the spike protein during fusion ^54^. Intriguingly, despite having different conformations resulting from the structural rearrangement during fusion, the epitope in pre- and post-fusion spike conformation could still be bound by both antibodies, suggesting either epitope recognition of the structurally conserved part of the epitope in the two spike conformations, or through an ‘induced fit’ mechanism at the binding interface ^55^.

Neutralization of authentic human-infecting coronaviruses, other than MERS-CoV, was not observed by these two antibodies, although weak cross-neutralizing capacity against pseudotyped viruses was seen for 28D9. This monoclonal antibody displayed high ELISA-reactivity to the spike proteins of HCoV-OC43, SARS-CoV, SARS-CoV-2. The lack of robust cross-neutralization by 28D9 may result from the approximately 5- to 10-fold higher equilibrium dissociation constants towards these spike proteins, relative to MERS-S, inferred by subtle differences in epitope composition. Structural insight into the 28D9 antibody binding mode to the stem helix epitope of targeted CoV spike proteins may guide antibody affinity improvement by structure-based design to increase breadth of neutralization.

The conserved stem helix epitope appears to be highly immunogenic. This is demonstrated by the independent isolation of three cross-reactive antibodies (28D9, 1.6C7 and 18H2) all targeting the stem-helix epitope from three independent mouse immunization experiments. In addition, antibodies towards this epitope are also elicited during natural infection, as demonstrated by the spike protein peptide microarray analysis using MERS-CoV-positive human and dromedary camel sera. Moreover, recent spike protein peptide microarray analyses by Li *et al.* ^56^ revealed that the SARS-CoV-2 peptide F^^1148^^K<u>EELDKYFK</u>NH^^1159^^ encompassing the 28D9/1.6C7 targeted stem helix epitope (underlined in sequence) was recognized by serum antibodies in ~90% of COVID-19 patients, whereas Ladner *et al.* ^57^ also identifies the stem-helix epitope region – with F^^1148^^K<u>EELDKYF</u>^^1156^^ as the minimal reactive SARS2-S peptide sequence - as the most widely-recognized SARS-CoV-2 linear epitope target in convalescent donors. It is tempting to speculate that the exceptionally high seroprevalence of stem-epitope targeting antibodies in COVID-19 patients is due to boosting of pre-existing immune response towards this conserved epitope resulting from earlier encounters with betacoronaviruses such as HCoV-OC43 and HCoV-HKU1 that are endemic in humans ^21^. Whether cross-reactive antibodies in humans towards the stem-helix or other conserved spike epitopes play a role in cross-protection or enhanced disease requires further investigation.

Together, the isolated cross-reactive antibodies define a conserved, immunogenic and vulnerabe site on the coronavirus spike protein. The discovery of eptiopes on viral glycoproteins targeted by cross-reactive neutralizing antibodies has fueled the design of broad range therapeutics and vaccines for other RNA viruses that display antigenic variation or show zoonotic potential, such as HIV-1 and influenza viruses ^13, 15, 58–60^. We made a first step into the identification of conserved sites among antigenically highly divergent coronavirus spike proteins. These efforts may enable the new generation of broadly protective vaccines and therapeutics, that can mitigate the potential risk of antigenic drift upon continuous circulation of coronaviruses in the population, as well as the looming threat of novel coronavirus emergence in humans.

## Methods

### Expression and purification of coronavirus spike proteins

Coronavirus spike ectodomains of MERS-CoV (residues 19–1262; strain EMC; GenBank accession number (GB): YP_009047204.1), SARS-CoV (residues 15–1182; strain Urbani; GB: AY278741.1), MHV (residues 15–1231; strain A59; UniProt accession number: P11224) and the S2 ectodomain of MHV (residues 718–1252; strain A59; UniProt accession number: P11224) fused with a C-terminal T4/GCN4 trimerization motif, a thrombin cleavage site and a Strep-tag purification tag were cloned in-frame into pMT\Bip\V5\His expression vector. The furin cleavage site at the S1/S2 junction was mutated to prevent cleavage by furin at this position. Spike ectodomains were stably produced in *Drosophila* S2 cell line, as previously described ^28^. Spike ectodomains of SARS-CoV-2 (residues 1–1,213; strain Wuhan-Hu-1; GenBank: QHD43416.1), HCoV-OC43 (residues 15–1263; strain Paris; UniProtKB: Q696P8) and HCoV-HKU1 (residues 14–1266; strain Caen1; GenBank: HM034837) were expressed transiently in HEK-293T cells with a C-terminal trimerization motif and Strep-tag using the pCAGGS expression plasmid. The genes encoding MERS-S2 ectodomain (residues 752–1262; strain EMC; GB: YP_009047204.1) were cloned in-frame between the HBM secretion signal peptide and a triple Strep-tag for purification in the pFastBac transfer vector. Generation of bacmid DNA and recombinant baculovirus was performed according to protocols from the Bac-to-Bac system (Invitrogen). All secreted proteins were purified from culture supernatant using streptactin beads (IBA) following the manufacturer’s protocol. All variants were generated using Q5^®^ High-Fidelity DNA Polymerase (NEB)-based site-directed mutagenesis.

### Generation of cross-reactive H2L2 mAbs

Six H2L2 mice were sequentially immunized in two weeks intervals with purified spike ectodomains of different CoVs in the following order ^34^: HCoV-OC43, SARS-CoV, MERS-CoV, HCoV-OC43, SARS-CoV and MERS-CoV. Antigens were injected at 20-25 μg/mouse using Stimune Adjuvant (Prionics) freshly prepared according to the manufacturer’s instruction for the first injection, while boosting was done using Ribi (Sigma) adjuvant. Injections were done subcutaneously into the left and right groin each (50 μl) and 100 μl intraperitoneally. Four days after the last injection, spleen and lymph nodes were harvested, and hybridomas made by standard method using SP 2/0 myeloma cell line (ATCC #CRL-1581) as a fusion partner. Hybridomas were screened in an antigen-specific ELISA and positives selected for further development, subcloned and produced on a small scale (100 ml of medium). For this purpose, hybridomas were cultured in serum- and protein-free medium for hybridoma culturing (PFHM-II (1X), Gibco) with addition of non-essential amino acids (100X NEAA, Biowhittaker Lonza, cat. no. BE13-114E). H2L2 antibodies were purified from hybridoma culture supernatants using Protein-G affinity chromatography. Purified antibodies were stored at 4°C for further use. DNA immunizations of H2L2 mice were done in two weekly intervals with spike protein expression plasmids in the following order: pCAGGS-OC43-S, pCAGGS-SARS-S and pCAGGS-MERS-S. During the procedure that involved shaving of the lower back, intradermal DNA injection (40-50 μg of DNA per mouse in 20-30 μl volume) and electroporation, mice were anesthetized. Electroporation was done according to instructions of manufacturer of the electroporation apparatus (Agile plus ID in vivo delivery system, BTX). In short, immediately after the DNA injection, mice were subjected to 10 electro pulses using 2×6 array needle surrounding the bleb (small blister under the skin) formed after the injection. Anaesthesia was given following standard operation procedures of the facility. Blood samples will be taken after the 4th DNA injectiion/electroporation. Five mice that developed satisfactory ELISA titres for all 3 antigens after DNA priming were additionally injected subcutaneously in two weeks intervals with 25 μg of each of the soluble trimeric spike proteins with Ribi adjuvant. 3-5 days after the last injection, mice were sacrificed and spleens and lymph nodes used to make a single cell suspension for the fusion experiment.

Animal studies were done under the animal permit AVD101002016512, under work protocol 16-512-22 called “heterologous prime-boost approach”, approved by the CCD (central committee for animal experiments).

### Production of recombinant human monoclonal antibodies

Production of recombinant human antibodies using HEK-293T was described previously ^43^. Briefly, the variable heavy (VH) and light (VL) chain sequences were amplified from cDNA and separately cloned into the expression plasmids with human IgG1 heavy chain and kappa chain constant regions, respectively (Invivogen). Both plasmids contain the interleukin-2 signal sequence to enable efficient secretion of recombinant antibodies. Recombinant human antibodies were expressed in HEK-293T cells following transient transfection with pairs of the IgG1 heavy and light chain expression plasmids according to protocols from Invivogen. Recombinant antibodies were purified using Protein A sepharose (IBA) according to the manufacturer’s instruction. mAb 1.6C7 and 7.7G6 used in the animal experiments were produced in Chinese hamster ovary (CHO) cells as previously described ^61^. The VH and VL sequences were synthesized by GeneArt, ThermoFisher and cloned into OriP-containing expression vectors, suitable for IgG1 production ^62^. A 20 L cell culture in a disposable rocking bioreactor was subsequently transiently transfected with the heavy and light chain expression vectors. The clarified harvest supernatant was purified using Protein A-based chromatography ^63^.

### ELISA analysis of antibody binding to CoV spike antigens

Purified coronavirus spike ectodomains were coated onto 96-well NUNC Maxisorp plates (Thermo Scientific) at equimolar amount at room temperature (RT) for 3 h followed by three washing steps with Phosphate Saline Buffer (PBS) containing 0.05% Tween-20. Plates were blocked with 5% milk (Protifar, Nutricia) in PBS with 0.1% Tween-20 at 4℃ overnight. Antibodies were allowed to bind to the plates at 4-fold serial dilutions, starting at 10 μg/ml diluted in PBS containing 3% BSA and 0.1% Tween20, at RT for 1 h. Antibody binding to the spike proteins was determined using a 1:2000 diluted HRP conjugated goat anti-human IgG (ITK Southern Biotech) for 1 h at RT and tetramethylbenzidine substrate (BioFX). Readout for binding was done at 450 nm (OD_450_) using the ELISA plate reader (EL-808, Biotek). Half-maximum effective concentration (EC_50_) binding values were calculated by 4-parameter logistic regression on the binding curves using GraphPad Prism version 7.04. To determine whether antibodies recognize a linear or conformational epitope, NUNC Maxisorp plates were coated with 100 ng/well of MERS-S ectodomain at RT for 3 h. Antigens were treated with or without 50 μl of denaturing buffer (200 mM DTT and 4% SDS in PBS) at 37℃ for 1 h. After three times washing steps with PBST (PBS with 0.05% Tween-20), plates were blocked by blocking buffer (5% milk in PBS with 0.1% Tween-20) at 4℃ overnight. Four-fold serially diluted primary antibodies were added to the plates and incubate at RT for 1 h. Plates were washed three times and antibody binding to the spike proteins was analysed as described above.

### Immunofluorescence microscopy

Antibody binding to cells expressing spike proteins of MERS, SARS-CoV, SARS-CoV-2, MHV, HCoV-OC43 and HCoV-HKU1 was measured by immunofluoresence microscopy. HEK-293T cells seeded on glass slides were transfected with plasmids encoding MERS-S, SARS-S, SARS2-S, MHV-S, HCoV-OC43-S or HCoV-HKU1-S C-terminally fused to the green fluorescence protein (GFP) using Lipofectamine 2000 (Invitrogen). Two days post transfection, cells were fixed by incubation with 2% paraformaldehyde in PBS for 20 min at RT before 0.1% Triton-100 permeabilization and stained for nuclei with 4,6-diamidino-2-phenylindole (DAPI). Cells were subsequently incubated with mAbs at a concentration of 10 μg/ml for 1 h at RT, followed by incubation with Alexa Fluor 594 conjugated goat anti-human IgG antibody (Invitrogen, Thermo Fisher Scientific) for 45 min at RT. The fluorescence images were recorded using a Leica SpeII confocal microscope.

### Flow cytometry

HEK-293T cells were seeded with a density of 3×10^6^ cells per T25 tissue culture flask. After reaching 70~80% confluency, cells were transfected with the pCAGGS expression plasmids encoding full-length spikes of MERS-CoV, SARS-CoV, SARS-CoV-2, MHV, HCoV-OC43 and HCoV-HKU1 C-terminally extended with GFP. Two days post transfection, cells were harvested by cell dissociation solution (Sigma-aldrich, Merck KGaA; cat. no. C5914). Single cell suspensions in FACS buffer (2% Fetal Bovine Serum (FBS), 5 mM EDTA and 0.02% NaN_3_ in PBS) were centrifuged at 400×g for 10 min. Cells were then treated with/without 0.1% Triton-100 after fixation in 3.7% paraformaldehyde. After a washing step in PBS, cells were blocked using 10% Normal Goat Serum (Gibco, Thermo Fisher Scientific, the Netherlands) diluted in PBS for 45 min at RT. Staining of spike proteins was performed by incubation of the cells with primary antibody (10 μg/ml) for 1 h at RT. They were then incubated with 1:200 diluted Alexa Fluor 594 conjugated goat anti-human IgG secondary antibody (Invitrogen, Thermo Fisher Scientific) for 45 min at RT and subjected to flow cytometric analysis with a CytoFLEX flow cytometer (Beckman). The results were analysed by FlowJo (version 10) and percentage of GFP^+^Alexa Fluor 594^+^ cells over GFP^+^ cells were calculated.

### Antibody binding kinetics and affinity measurement

The measurement of binding kinetics and affinity of antibodies to CoV spike ectodomains was performed using biolayer interferometry (Octet RED384 machine) as described before ^43^. Briefly, fully human antibodies with optimal concentration (44 nM) which showed the desired loading curve characteristics and high signal in the association step were loaded onto Protein A biosensors for 10 min. Binding of CoV spikes was performed by incubating the biosensor with various concentrations of recombinant spike ectodomain (1600-800-400-200-100-50-25-12.5-6.25 nM) for 10 min followed by the dissociation step which was run long enough (60 min) to observe the decrease of the binding response. The affinity constant *K_D_* was calculated using 1:1 Langmuir binding model on Fortebio Data Analysis 7.0 software.

### Pseudotyped virus neutralization assay

The production of coronavirus spike pseudotyped VSV virus and the neutralization assay was performed as described previously ^43, 64^. In brief, HEK-293T cells at 70~80% confluency were transfected with the pCAGGS expression vectors encoding full-length MERS-S, SARS-S, SARS2-S or OC43-S with a C-terminal cytoplasmic tail truncation to increase cell surface expression levels. In case of OC43-S, cells were co-transfected with pCAGGS vector encoding the Fc-tagged bovine coronavirus hemagglutinin esterase (HE-Fc) protein at molar ratios of 8:1 (S:HE-Fc). Forty-eight hours post transfection, cells were infected with VSV-G pseudotyped VSVΔG bearing the firefly (*Photinus pyralis*) luciferase reporter gene at a MOI of 1. Twenty-four hours later, supernatant was harvested and filtered through 0.45 μm membrane. Pseudotyped VSV virus were titrated on monolayer African green monkey kidney VeroCCL81 cells (MERS-S pseudotyped VSV), VeroE6 cells (SARS-S and SARS2-S pseudotyped VSV) or on HRT-18 cells (OC43-S pseudotyped VSV). In the virus neutralization assay, serially diluted mAbs were pre-incubated with an equal volume of virus at RT for 1 h, and then inoculated on Vero/HRT-18 cells, and further incubated at 37℃. After 20 h, cells were washed once with PBS and lysed with cell lysis buffer (Promega). The expression of firefly luciferase was measured on a Berthold Centro LB 960 plate luminometer using D-luciferin as a substrate (Promega). The percentage of infectivity was calculated as the ratio of luciferase readout in the presence of mAbs normalized to luciferase readout in the absence of mAb. The half maximal inhibitory concentrations (IC_50_) were determined using 4-parameter logistic regression (GraphPad Prism v7.0).

### Authentic virus neutralization assay

Neutralization of authentic MERS-CoV, SARS-CoV and SARS-CoV-2 was performed using a plaque reduction neutralization test (PRNT) as described earlier, with some modifications ^65, 66^. In brief, mAbs were serially diluted and mixed with MERS-CoV, SARS-CoV or SARS-CoV-2 for 1 hour. The mixture was then added to Huh-7 cells (MERS-CoV) or VeroE6 cells (SARS-CoV and SARS-CoV-2) and incubated for 1 hr, after which the cells were washed and further incubated in medium for 8 h. Subsequently, the cells were washed, fixed, permeabilized and the infection was detected using immunofluorescent staining using antibodies specific for the viruses used. The signal was developed using a precipitate forming TMB substrate (True Blue, KPL) and the number of infected cells per well were counted using the ImmunoSpot® Image analyzer (CTL Europe GmbH). The half maximal inhibitory concentrations (IC_50_) were determined using 4-parameter logistic regression (GraphPad Prism version 8).

### Receptor binding inhibition assay

The DPP4 receptor binding inhibition assay was performed as described previously ^43^. Recombinant soluble DPP4 was coated on NUNC Maxisorp plates (Thermo Scientific) at 100 ng/well at RT for 3 h. Plates were washed three times with PBS containing 0.05% Tween-20 and blocked with 5% milk (Protifar, Nutricia) in PBS containing 0.1% Tween-20 at 4℃ overnight. Recombinant MERS-CoV S ectodomain and serially diluted mAbs were mixed and incubated for 1 h at RT. The mixture was added to the plate for 1 h at RT, after which plates were washed three times. Binding of MERS-CoV S ectodomain to DPP4 was detected using 1:1000 diluted HRP-conjugated anti-StrepMAb (IBA) that recognizes the Strep-tag affinity tag on the MERS-CoV S ectodomain. Detection of HRP activity was performed as described above (ELISA section).

### Fusion inhibition assay

Fusion inhibition assay was perfomed as described ^43^, with some adaptations. Huh-7 cells were seeded one day before reaching a confluency of 70-80%. Cells were transfected with pCAGGS expression plasmid encoding full-length MERS-S C-terminally fused with a GFP-tag using Lipofectamine 2000. The furin cleavage site *R*^747^SVR^751^ at S1/S2 junction was mutated to *K*SVR to avoid the cleavage by endogenous proteases. At 48 h post transfection, cells were pre-treated with DMEM only or DMEM with 20 μg/ml mAbs for 1 h and subsequently treated with DMEM with 20 μg/ml of exogenous trypsin to activate the spike fusion function at 37°C for 2 h. Cells were fixed with 3.7% paraformaldehyde after observation of the syncytia formation. 4,6-diamidino-2-phenylindole (DAPI) was used to stain the nuclei. The expression of MERS-S was confirmed based on the GFP signal, and the cell-cell fusion was monitored by large GFP-fluorescent muti-nucleated syncytia. The fluorescence images were recorded using the EVOS FL fluorescence microscope (Thermo Fisher Scientific, the Netherlands).

### Antibody binding competition assay

Antibody binding competition was performed using biolayer interferometry (Octet Green; ForteBio), as described previously ^43^. In brief, MERS-CoV spike antigen 50 μg/ml was immobilized onto the anti-strep mAb-coated protein A biosensor. After a brief washing step, the biosensor tips were immersed into a well containing primary mAb at a concentration of 50 μg/ml for 15 min and subsequently into a well containing the competing mAb (secondary mAb) at a concentration of 50 μg/ml for 15 min. A 5-min washing step in PBS was included in between steps.

### Spike protein peptide microarray analysis

Spike protein peptide microarray analysis to map linear epitopes of antibodies in convalescent sera was performed by PEPSCAN (Lelystad, The Netherlands). Overlapping peptides that cover the entire MERS-CoV spike ectodomain (residues 1-1,296) were synthesized with an offset of one or two residues. Order of these peptides was randomized, when synthesized on mini-cards. ELISA reactivity of 5 human (H1 to H5) and 4 dromedary camel (D1 to D4) MERS-positive sera was assessed, as well as a MERS-negative serum from human (H-CTRL) or camel (D-CTRL). The binding of antibody to each of the synthesized peptides was tested in a PEPSCAN-based ELISA. The peptide arrays were incubated with the primary antibody solution (overnight at 4°C). After washing, the peptide arrays were incubated with a 1/1000 dilution of an appropriate antibody peroxidase conjugate (goat anti-human HRP conjugate, Southern Biotech, cat. no.: 2010-05 or goat anti-lama HRP conjugate, Abcore, cat. no. AC15-0354) for 1 h at 25°C. After washing, the peroxidase substrate 2,2’-azino-di-3-ethylbenzthiazoline sulfonate (ABTS) and 20 μl/ml of 3% H2O2 was added. After 1 h, the color development was measured and quantified with a charge coupled device (CCD) - camera and an image processing system. The values obtained from the CCD camera range from 0 to 3000 mAU. Samples were scaled per serum sample using a cut-off of twice the mean absorbance obtained for each serum. The use of human materials was approved by the local medical ethical committee (MEC approval: 2014-414).

To map the epitopes of the monoclonal antibodies, 30-amino acid long peptides (with 15-a.a. overlap) were synthesized (Genscript) covering the conserved C-terminal part of the MERS-S2 ectodomain (residues 869-1,288). 100 ng/well of each peptide was coated onto 96-well NUNC Maxisorb plate at 4°C overnight. Followed by three washing steps with PBST (PBS with 0.05% Tween-20), plates were blocked with 5% milk (Protifar, Nutricia) in PBS with 0.1% Tween-20 at RT for 3 h. Antibodies were allowed to bind to the plates at 4-fold serial dilutions, starting at 10 μg/ml, at RT for 1 h. Antibody binding to the peptides was determined using a goat anti-human IgG HRP conjugate (ITK Southern Biotech) for 1 h at RT and tetramethylbenzidine substrate (BioFX). Readout for binding was done as described in the ELISA section. To identify critical residues for mAb binding, a single alanine mutation was introduced on the 15-mer spike peptide fragment that comprises the linear epitope. Reactivity of antibodies with peptides with a single alanine substitution was measured by ELISA according to the method described above.

### Passive immunization and protection tests of mice

*In vivo* prophylactic and therapeutic efficacy of 1.6C7 against MERS-CoV infection was evaluated in the transgenic mouse model K18 TghDpp4 expressing the receptor for the human MERS-CoV ^46^. Groups of 8 mice, 20-30 weeks old, were given 50 μg of 1.6C7 (equivalent to 1.8 mg of the antibody per kg) by intraperitoneal injection, 24 hours before or after intranasal infection with a lethal dose of MERS-CoV (EMC isolate; 5 × 10^3^ PFU/mouse). The potent neutralizing anti-MERS-S1 control antibody 7.7G6 ^43^ and an isotype matched negative control mAb were taken along as controls. In a second experiment, the prophylactic efficacy of mAb 28D9 was tested against MERS-CoV and SARS-CoV infection in the K18 TghDpp4 mouse model. Groups of 16-20-week old mice (n=5), were given 50 or 200 μg 28D9 or isotype control antibody (equivalent to 1.8 or 7.2 mg of the antibody per kg, respectively) by intraperitoneal injection, 24 h before intranasal infection with a lethal dose of MERS-CoV (EMC isolate; 5 × 10^3^ PFU/mouse) or mouse adapted SARS-CoV (strain SARS-CoV-MA15-WT-M2; 1 × 10^5^ PFU/mouse). Animal protection studies were done under the animal permit PROEX-199/19, approved by the Community of Madrid (Spain), and performed in biosafety level 3 facilities at CISA-INIA (Madrid).

MERS-CoV titers and lung histopathology were tested as described earlier ^65^. To analyze MERS-CoV titers, one fourth of the right lung was homogenized using a MACS homogenizer (Miltenyi Biotec) according to manufacturer’s protocols. Virus titrations were performed on Huh-7 cells following standard procedures. In brief, cells were overlaid with DMEM containing 0.6% low-melting agarose and 2% FBS, fixed with 10% formaldehyde and stained with 0.1% crystal violet at 72 h post infection. The left lung of infected mice was fixed in 10% zinc formalin for 24 h at 4°C and paraffin embedded for lung histopathological examination. Serial longitudinal 5 μm-sections of formaling were stained with hematoxylin and eosin (H&E) and subjected to histopathological examination with a ZEISS Axiophot fluorescence microscope. Samples were obtained using a systematic uniform random procedure, consisting in serial parallel slices made at a constant thickness interval of 50 μm.

## Acknowledgements

We thank dr. Yoshiharu Matsuura (Osaka University, Japan) for providing the luciferase-encoding VSV-G pseudotyped VSVΔG-luc virus, Volker Thiel (University of Bern, Switzerland) for providing the HCoV-HKU1 spike protein encoding plasmid and Alejandra Tortorici (Institute Pasteur, France) for providing MERS-CoV S ectodomain. We thank Jaap Willem Back from PEPSCAN Presto BV, Lelystad, the Netherlands for his assistance with the spike protein peptide microarray analysis. We thank Ludo Broos for technical support. This study was done within the framework of the National Centre for One Health (NCOH) and the Utrecht Molecular Immunology Hub - Utrecht University. Funding: The project was co-financed by a grant from the Zoonotic Anticipation and Preparedness Initiative [ZAPI project; Innovative Medicines Initiative (IMI) grant agreement no. 115760], with the assistance and financial support of IMI and the European Commission, and in-kind contributions from European Federation of Pharmaceutical Industries and Associations partners. The collaboration project is co-funded by the PPP Allowance made available by Health~Holland, Top S_ecto_r Life Sciences & Health, to stimulate public-private partnerships. This study was also partially financed by grants from the Ministry of Science and Innovation of Spain (BIO2016-75549-R AEI/FEDER, UE) and NIH (2PO1AIO6O699). The mice used to generate the mAbs produced in this study were provided by Harbour Antibodies BV, a daughter company of Harbour Biomed (http://www.harbourbiomed.com). Chunyan Wang was supported by a grant from the China Scholarship Council.

## Author Contributions

B.J.B. conceived and coordinated the study. C.W. D.D. and B.J.B. designed the experiments. C.W., R.H., J.G.A., W.L., N.M.A.O., I.A., I.W., B.D., R.F.D., I.S. and D.D. conducted the experiments. D.L.H., O.D., F.G., F.J.M.K., B.L.H., L.E., D.D. and B.J.B. supervised part of the experiments. All authors contributed to the interpretations and conclusions presented. C.W. and B.J.B. wrote the manuscript with comments from all co-authors. All authors participated in editing the manuscript.

## Competing interests

C.W., R.v.H., W.L., I.W., B.v.D., N.M.A.O., F.G., F.J.M.v.K., B.L.H., D.D., and B.J.B. are inventors on a patent application on monoclonal antibodies targeting MERS-CoV (patent publication no.: WO/2020/169755). F.G., D.D. and R.H. are non-substantial interest shareholders in Harbour Biomed and were part of the team that generated the mice.

## Data availability

All data are available from the corresponding author upon reasonable requests.

## References

1. Zhou, P. et al. A pneumonia outbreak associated with a new coronavirus of probable bat origin. Nature 579, 270–273 (2020).

2. Gorbalenya, A. et al. Coronaviridae Study Group of the International Committee on Taxonomy of Viruses. The species severe acute respiratory syndrome-related coronavirus: classifying 2019-nCoV and naming it SARS-CoV-2. Nature microbiology 2020, 03–04 (2020).

3. Peiris, J., Guan, Y. & Yuen, K. Severe acute respiratory syndrome. Nat. Med. 10, S88–S97 (2004).

4. World Health Organization. MERS situation update, January 2020. URL: http://www.emro.who.int/health-topics/mers-cov/mersoutbreaks.html (29.02.2020) (2020).

5. Gaunt, E. R., Hardie, A., Claas, E. C., Simmonds, P. & Templeton, K. E. Epidemiology and clinical presentations of the four human coronaviruses 229E, HKU1, NL63, and OC43 detected over 3 years using a novel multiplex real-time PCR method. J. Clin. Microbiol. 48, 2940–2947 (2010).

6. Lim, Y. X., Ng, Y. L., Tam, J. P. & Liu, D. X. Human coronaviruses: a review of virus– host interactions. Diseases 4, 26 (2016).

7. Walsh, E. E., Shin, J. H. & Falsey, A. R. Clinical impact of human coronaviruses 229E and OC43 infection in diverse adult populations. J. Infect. Dis. 208, 1634–1642 (2013).

8. Morfopoulou, S. et al. Human coronavirus OC43 associated with fatal encephalitis. N. Engl. J. Med. 375, 497–498 (2016).

9. Su, S. et al. Epidemiology, genetic recombination, and pathogenesis of coronaviruses. Trends Microbiol. 24, 490–502 (2016).

10. Chan, J. F., To, K. K., Tse, H., Jin, D. & Yuen, K. Interspecies transmission and emergence of novel viruses: lessons from bats and birds. Trends Microbiol. 21, 544–555 (2013).

11. Hu, B., Ge, X., Wang, L. & Shi, Z. Bat origin of human coronaviruses. Virology journal 12, 1–10 (2015).

12. Burton, D. R. & Walker, L. M. Rational vaccine design in the time of COVID-19. Cell Host & Microbe 27, 695–698 (2020).

13. Burton, D. R., Poignard, P., Stanfield, R. L. & Wilson, I. A. Broadly neutralizing antibodies present new prospects to counter highly antigenically diverse viruses. Science 337, 183–186 (2012).

14. Pappas, L. et al. Rapid development of broadly influenza neutralizing antibodies through redundant mutations. Nature 516, 418–422 (2014).

15. Chen, Y. et al. Influenza infection in humans induces broadly cross-reactive and protective neuraminidase-reactive antibodies. Cell 173, 417–429. e10 (2018).

16. Flyak, A. I. et al. Cross-reactive and potent neutralizing antibody responses in human survivors of natural ebolavirus infection. Cell 164, 392–405 (2016).

17. Wec, A. Z. et al. Antibodies from a human survivor define sites of vulnerability for broad protection against ebolaviruses. Cell 169, 878–890. e15 (2017).

18. Gilchuk, P. et al. Multifunctional pan-ebolavirus antibody recognizes a site of broad vulnerability on the ebolavirus glycoprotein. Immunity 49, 363–374. e10 (2018).

19. Chan, K. et al. Cross-reactive antibodies in convalescent SARS patients’ sera against the emerging novel human coronavirus EMC (2012) by both immunofluorescent and neutralizing antibody tests. J. Infect. 67, 130–140 (2013).

20. Barnes, C. O. et al. Structures of human antibodies bound to SARS-CoV-2 spike reveal common epitopes and recurrent features of antibodies. bioRxiv (2020).

21. Wec, A. Z. et al. Broad neutralization of SARS-related viruses by human monoclonal antibodies. Science 369, 731–736 (2020).

22. Kirchdoerfer, R. N. et al. Stabilized coronavirus spikes are resistant to conformational changes induced by receptor recognition or proteolysis. Scientific reports 8, 1–11 (2018).

23. Li, Z. et al. The human coronavirus HCoV-229E S-protein structure and receptor binding. Elife 8, e51230 (2019).

24. Park, Y. et al. Structures of MERS-CoV spike glycoprotein in complex with sialoside attachment receptors. Nature structural & molecular biology 26, 1151–1157 (2019).

25. Shang, J. et al. Cryo-EM structure of infectious bronchitis coronavirus spike protein reveals structural and functional evolution of coronavirus spike proteins. PLoS pathogens 14, e1007009 (2018).

26. Shang, J. et al. Cryo-Electron Microscopy Structure of Porcine Deltacoronavirus Spike Protein in the Prefusion State. J. Virol. 92, 10.1128/JVI.01556-17 (2018).

27. Tortorici, M. A. et al. Structural basis for human coronavirus attachment to sialic acid receptors. Nature structural & molecular biology 26, 481–489 (2019).

28. Walls, A. C. et al. Cryo-electron microscopy structure of a coronavirus spike glycoprotein trimer. Nature 531, 114–117 (2016).

29. Walls, A. C. et al. Unexpected receptor functional mimicry elucidates activation of coronavirus fusion. Cell 176, 1026–1039. e15 (2019).

30. Yuan, Y. et al. Cryo-EM structures of MERS-CoV and SARS-CoV spike glycoproteins reveal the dynamic receptor binding domains. Nature communications 8, 1–9 (2017).

31. Cai, Y. et al. Distinct conformational states of SARS-CoV-2 spike protein. bioRxiv (2020).

32. Wrapp, D. et al. Cryo-EM structure of the 2019-nCoV spike in the prefusion conformation. Science 367, 1260–1263 (2020).

33. Menachery, V. D. et al. A SARS-like cluster of circulating bat coronaviruses shows potential for human emergence. Nat. Med. 21, 1508–1513 (2015).

34. Wang, C. et al. A human monoclonal antibody blocking SARS-CoV-2 infection. Nature communications 11, 1–6 (2020).

35. Zhou, D. et al. Structural basis for the neutralization of SARS-CoV-2 by an antibody from a convalescent patient. Nature Structural & Molecular Biology, 1–9 (2020).

36. Brouwer, P. et al. Potent neutralizing antibodies from COVID-19 patients define multiple targets of vulnerability. bioRxiv (2020).

37. Lv, H. et al. Cross-reactive antibody response between SARS-CoV-2 and SARS-CoV infections. Cell Reports, 107725 (2020).

38. Pinto, D. et al. Cross-neutralization of SARS-CoV-2 by a human monoclonal SARS-CoV antibody. Nature, 1–6 (2020).

39. Lv, Z. et al. Structural basis for neutralization of SARS-CoV-2 and SARS-CoV by a potent therapeutic antibody. Science 369, 1505–1509 (2020).

40. Rogers, T. F. et al. Isolation of potent SARS-CoV-2 neutralizing antibodies and protection from disease in a small animal model. Science 369, 956–963 (2020).

41. Haagmans, B. L. et al. SARS-CoV-2 neutralizing human antibodies protect against lower respiratory tract disease in a hamster model. bioRxiv (2020).

42. Walls, A. C. et al. Structure, function, and antigenicity of the SARS-CoV-2 spike glycoprotein. Cell (2020).

43. Widjaja, I. et al. Towards a solution to MERS: protective human monoclonal antibodies targeting different domains and functions of the MERS-coronavirus spike glycoprotein. Emerging microbes & infections 8, 516–530 (2019).

44. Walls, A. C. et al. Glycan shield and epitope masking of a coronavirus spike protein observed by cryo-electron microscopy. Nature structural & molecular biology 23, 899 (2016).

45. Walls, A. C. et al. Tectonic conformational changes of a coronavirus spike glycoprotein promote membrane fusion. Proc. Natl. Acad. Sci. U. S. A. 114, 11157–11162 (2017).

46. Li, K. et al. Middle East respiratory syndrome coronavirus causes multiple organ damage and lethal disease in mice transgenic for human dipeptidyl peptidase 4. J. Infect. Dis. 213, 712–722 (2016).

47. Ekiert, D. C. et al. Cross-neutralization of influenza A viruses mediated by a single antibody loop. Nature 489, 526–532 (2012).

48. Schmidt, A. G. et al. Viral receptor-binding site antibodies with diverse germline origins. Cell 161, 1026–1034 (2015).

49. Routledge, E., Stauber, R., Pfleiderer, M. & Siddell, S. G. Analysis of murine coronavirus surface glycoprotein functions by using monoclonal antibodies. J. Virol. 65, 254–262 (1991).

50. Elshabrawy, H. A., Coughlin, M. M., Baker, S. C. & Prabhakar, B. S. Human monoclonal antibodies against highly conserved HR1 and HR2 domains of the SARS-CoV spike protein are more broadly neutralizing. Plos one 7, e50366 (2012).

51. Lip, K. M. et al. Monoclonal antibodies targeting the HR2 domain and the region immediately upstream of the HR2 of the S protein neutralize in vitro infection of severe acute respiratory syndrome coronavirus. J. Virol. 80, 941–950 (2006).

52. Supekar, V. M. et al. Structure of a proteolytically resistant core from the severe acute respiratory syndrome coronavirus S2 fusion protein. Proc. Natl. Acad. Sci. U. S. A. 101, 17958–17963 (2004).

53. Xu, Y. et al. Structural basis for coronavirus-mediated membrane fusion. Crystal structure of mouse hepatitis virus spike protein fusion core. J. Biol. Chem. 279, 30514–30522 (2004).

54. King, L. B. et al. Cross-reactive neutralizing human survivor monoclonal antibody BDBV223 targets the ebolavirus stalk. Nature communications 10, 1–8 (2019).

55. Berger, C. et al. Antigen recognition by conformational selection. FEBS Lett. 450, 149–153 (1999).

56. Li, Y. et al. Linear epitopes of SARS-CoV-2 spike protein elicit neutralizing antibodies in COVID-19 patients. medRxiv (2020).

57. Ladner, J. T. et al. Epitope-resolved profiling of the SARS-CoV-2 antibody response identifies cross-reactivity with an endemic human CoV. bioRxiv (2020).

58. Xu, K. et al. Epitope-based vaccine design yields fusion peptide-directed antibodies that neutralize diverse strains of HIV-1. Nat. Med. 24, 857–867 (2018).

59. Nachbagauer, R. & Krammer, F. Universal influenza virus vaccines and therapeutic antibodies. Clinical Microbiology and Infection 23, 222–228 (2017).

60. Crowe Jr, J. E. Principles of broad and potent antiviral human antibodies: insights for vaccine design. Cell host & microbe 22, 193–206 (2017).

61. Daramola, O. et al. A high-yielding CHO transient system: coexpression of genes encoding EBNA-1 and GS enhances transient protein expression. Biotechnol. Prog. 30, 132–141 (2014).

62. Gahn, T. A. & Sugden, B. An EBNA-1-dependent enhancer acts from a distance of 10 kilobase pairs to increase expression of the Epstein-Barr virus LMP gene. J. Virol. 69, 2633–2636 (1995).

63. Liu, H. F., Ma, J., Winter, C. & Bayer, R. Recovery and purification process development for monoclonal antibody production (MAbs Ser. 2, Taylor & Francis, 2010).

64. Hulswit, R. J. G. et al. Human coronaviruses OC43 and HKU1 bind to 9-O-acetylated sialic acids via a conserved receptor-binding site in spike protein domain A. Proc. Natl. Acad. Sci. U. S. A. 116, 2681–2690 (2019).

65. Raj, V. S. et al. Chimeric camel/human heavy-chain antibodies protect against MERS-CoV infection. Science advances 4, eaas9667 (2018).

66. Okba, N. M. et al. Severe acute respiratory syndrome coronavirus 2− specific antibody responses in coronavirus disease patients. Emerging infectious diseases 26, 1478–1488 (2020).

67. Madeira, F. et al. The EMBL-EBI search and sequence analysis tools APIs in 2019. Nucleic Acids Res. 47, W636–W641 (2019).

68. Pettersen, E. F. et al. UCSF Chimera—a visualization system for exploratory research and analysis. Journal of computational chemistry 25, 1605–1612 (2004).

